# Larixol is not an inhibitor of Gα_i_ containing G proteins and lacks effect on signaling mediated by human neutrophil expressed formyl peptide receptors

**DOI:** 10.1101/2023.10.11.561907

**Authors:** Lena Björkman, Huamei Forsman, Linda Bergqvist, Claes Dahlgren, Martina Sundqvist

## Abstract

Neutrophils express several G protein-coupled receptors (GPCRs) connected to intracellular Gα_i_ or Gα_q_ containing G proteins for down-stream signaling. To dampen GPCR mediated inflammatory processes, several inhibitors targeting the receptors and/or their down-stream signals, have been developed. Potent and selective inhibitors for Gα_q_ containing G proteins are available, but potent and specific inhibitors of Gα_i_ containing G proteins are lacking. Recently, Larixol, a compound extracted from the root of *Euphorbia formosana*, was shown to abolish human neutrophil functions induced by *N*-formyl-methionyl-leucyl-phenylalanine (fMLF), an agonist recognized by formyl peptide receptor 1 (FPR1) which couple to Gα_i_ containing G proteins. The inhibitory effect was suggested to be due to interference with/inhibition of signals transmitted by βγ complexes of the Gα_i_ containing G proteins coupled to FPR1. In this study, we applied Larixol, obtained from two different commercial sources, to determine the receptor- and G protein-selectivity of this compound in human neutrophils. However, our data show that Larixol not only lacks inhibitory effect on neutrophil responses mediated through FPR1, but also on responses mediated through FPR2, a Gα_i_ coupled GPCR closely related to FPR1. Furthermore, Larixol did not display any features as a selective inhibitor of neutrophil responses mediated through the Gα_q_ coupled GPCRs for platelet activating factor and ATP. Hence, our results imply that the inhibitory effects described for the root extract of *Euphorbia formosana* are not mediated by Larixol and that the search for a selective inhibitor of G protein dependent signals generated by Gα_i_ coupled neutrophil GPCRs must continue.

## 1. Introduction

G protein-coupled receptors (GPCRs) comprise the largest family of plasma membrane receptors in the human genome and include more than 800 different receptors [1, 2]. To date, around 35% of all drugs approved by the US Food and Drug Administration (FDA) act through GPCRs and ligands targeting GPCRs as well as reagents that regulate GPCR mediated responses are extensively exploited as novel drug candidates for various diseases [3–5]. All members of the GPCR family have seven transmembrane-spanning α-helices that connect three intracellular loops and the C-terminal tail to three extracellular loops and the N-termini [6]. The cellular activities regulated by the GPCRs are initiated by binding of an agonist to the so called orthosteric recognition cavity exposed on the surface of the cells expressing the receptor, and this binding structurally changes the receptor parts that couple to a signaling heterotrimeric G protein localized on the intracellular side of the membrane. The signal transduction upon agonist binding to the GPCRs is initiated by an activation of the heterotrimeric G protein complex (comprising two subunits, an α subunit and a heterodimeric βγ subunit) and a dissociation of the GTP/GDP binding α subunit from β/γ protein dimer [7]. The human genome encodes for several different G protein subunits [8] comprising 16 different α subtypes [9], five different β subtypes and 12 different γ subtypes [10] which basically all can couple to each other leading to a large variety of the heterotrimeric G protein complex. Lately, the functional outcome of an activated GPCR based on its coupling to diverse β and γ subunits (which are tightly associated and form a β/γ protein dimer) has started to gain attention [10, 11]. However, the GPCRs are primarily classified based on their coupling to the more extensively studied α subunits, which based on subunit similarities classifies GPCRs as coupled to either a Gα_s_, Gα_12/13_, Gα_q_ or Gα_i_ containing heterotrimeric G protein [9]. Yet, it should be mentioned that most data that has defined a specific GPCRs’ coupling to either of the four classes of the heterotrimeric G proteins have been performed using cells that overexpress an exclusive GPCR that is combined with distinctive G proteins. This is primarily due to that the different signaling G proteins are very similar which makes it hard to perform such studies in naïve cells. In addition, the signaling similarity between different G proteins has also made it difficult to design/find selective inhibitors for a specific G protein which has led to that the identification of the signaling G protein coupled to a GPCRs expressed in naïve cells have been (and still is) challenging.

The available tool compounds used to identify the signaling G protein in naïve cells have for long been the toxins from the bacteria termed *Vibro cholera* (cholera toxin; inhibits Gα_s_ containing G proteins) and *Bordetella pertussis* (pertussis toxin; inhibits Gα_i_ containing G proteins). However, even though pertussis toxin is the best-established tool to inhibit Gα_i_ containing G proteins to date [9], it has some limitations including some Gα_i_ subunit independent effects in a variety of cells [12], requirement of long incubation times (≥ 1.5 hours) with human neutrophils for the inhibition to occur and doubtful specificity for inhibition of only the Gα_i_ subunit and not the Gα_q_ subunit regarding GPCRs expressed by human neutrophils [13, 14]. Novel, selective, and potent G protein inhibitors have thus been eagerly awaited and among the different G proteins, specific Gα_q_ depsipeptide inhibitors are now available [15, 16]. These have successfully been used to characterize G protein profiles for many GPCRs expressed in several different cells including human neutrophils. Based on the Gα_q_ depsipeptide inhibitors it has become obvious that some neutrophil expressed GPCRs such as the formyl peptide receptors (FPRs) are, as expected, insensitive to Gα_q_ inhibition whereas signaling through other receptors such as the PAF receptor (PAFR, the receptor for platelet activating factor [PAF]) and the purinergic receptor P2Y_2_ (P2Y_2_R, the receptor for ATP), are hindered by Gα_q_ inhibition, strongly suggesting that they couple to Gα_q_ containing G proteins [7]. Recently, a diterpene extract from the root of plant termed *Euphorbia formosana* was shown to inhibit neutrophil functions induced by the prototype neutrophil chemotactic peptide *N*-formyl-methionyl-leucyl-phenylalanine (fMLF), recognized by formyl peptide receptor 1 (FPR1). The active compound in the inhibiting root extract was identified as Larixol (chemical structure shown in Fig 1) and the inhibition was suggested to be achieved through an effect on the interaction between the inhibitor and the heterodimeric βγ protein dimer of the Gα_i_ containing G protein coupled to FPR1. Accordingly, the βγ subunit inhibitor present in the root of *Euphorbia formosana*, was shown to inhibit various fMLF mediated human neutrophil functions including the FPR1 mediated rise in the intracellular concentration of free calcium ions ([Ca^2+^]_i_) and activation of the superoxide anion generating NADPH-oxidase [17]. However, as no data on the receptor/G protein selectivity were presented in their study, we aimed to characterize the inhibitory effects of Larixol on human neutrophil GPCR signaling by including Gα_i_ coupled FPR1 and the closely related FPR2 [18, 19]), as well as the two Gα_q_ coupled GPCRs (the PAFR and P2Y_2_R [7]. Accordingly, we determined the inhibitory effects of Larixol on the response induced in human neutrophils by agonists specific for these GPCRs with focus on the agonist induced rise in [Ca^2+^]_i_ and the production of superoxide anions generated by the NADPH-oxidase. However, our results do not reveal any GPCR/G protein selectivity of Larixol or display any evidence that the compound is an inhibitor of fMLF-mediated neutrophil functions as the presence of Larixol (obtained from two different commercial sources) did not display any inhibitory effect on human neutrophil responses mediated by activation of either of the two Gα_i_ coupled FPRs (FPR1 or FPR2) expressed in human neutrophils.

**Figure 1.**
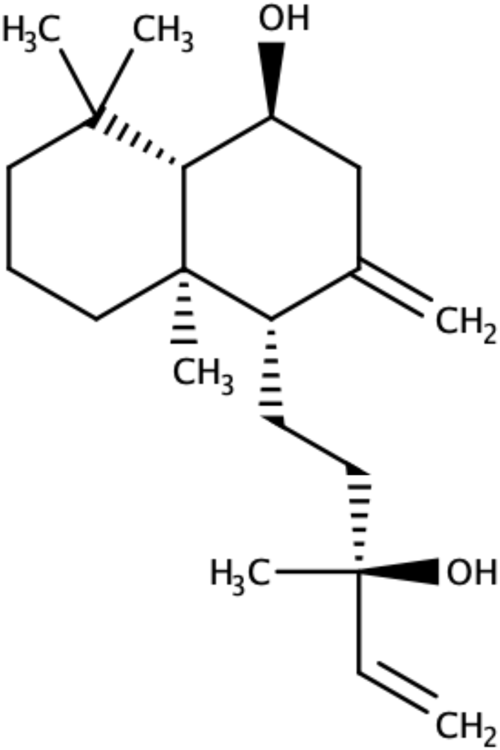
Chemical structure of Larixol.

## 2. Materials and methods

### 2.1 Chemicals and reagents

Cytiva Ficoll-Paque^TM^ Plus Media (#17-1440-03) was purchased from Fischer Scientific (Gothenburg, Sweden) and RPMI 1640 medium without phenol red (#11835030) was from Gibco (Thermo Fischer Scientific, Gothenburg, Sweden). Dextran T500 (#551005009007) was from Pharmacosmos (Holbaek, Denmark) and Fura-2-acetoxymethyl ester (AM, #F1221) was from Invitrogen by Thermo Fischer Scientific (Gothenburg, Sweden). Isoluminol (# A-8264), dimethyl sulfoxide (DMSO, #D2650), Latrunculin A (#L5163) and PMA (phorbol 12-myristate 13-acetate, #P8139) were from Sigma-Aldrich (Merck, Burlington, MA, USA) whereas horseradish peroxidase (HRP, peroxidase [POD], #10108090001) and bovine serum albumin fraction V (BSA, #10735094001) were from Roche (Merck, Burlington, MA, USA). The FPR1 agonist fMLF (*N*-formyl-methionyl-leucyl-phenylalanine, #F3506) and the P2Y_2_R agonists ATP (adenosine 5’-thriphosphate disodium salt hydrate, #A7699) and ATPγs (adenosine 5′-[γ-thio]triphosphate tetralithium salt #A1388) were also from Sigma-Aldrich whereas PAF (platelet activating factor, #84009) was from Avanti Polar Lipids (Birmingham, AL, USA) and the FPR2 agonist WKYMVM (tryptophyl-lysyl-tyrosyl-methinyl-valinyl-methioninamide) was synthesized by Caslo Aps (Lyngby, Denmark). The Gα_q_ inhibitor YM-254890 (#257-00631) was purchased from FUJIFILM Wako Chemicals Europe (Nordic Biolabs, Täby, Sweden). Larixol was from two different sources; 95% pure synthetic Larixol (#FL65054) was purchased from Biosynth (Staad, Switzerland) and 95% pure Larixol isolated from turpentine oil of *Larix sp* (#L175950) was obtained from Toronto research chemicals (Toronto, ON, Canada). All ligand stocks were dissolved, aliquoted and stored as follows; fMLF and WKYMVM (first dissolved to 10 mM in DMSO and then further diluted to 0.1 mM in dH_2_O, storage -80°C), PAF (dissolved to 1 mM in DMSO, storage -80°C), ATP and ATPγs (dissolved to 50 mM in dH_2_O, storage -20°C), YM-254890 (dissolved to 1 mM in DMSO, storage -20°C) and Larixol (dissolved to 100 mM in DMSO, storage -80°C). Subsequent dilutions of all ligands and reagents were made in Krebs-Ringer Glucose phosphate buffer (KRG; 120 mM NaCl, 4.9 mM KCl, 1.7 mM KH_2_PO_4_, 8.3 mM NaH_2_PO_4_, 1.2 mM MgSO_4_, 10 mM glucose, and 1 mM CaCl_2_ in dH_2_O, pH 7.3).

### 2.2. Ethics Statement

This study comprises leukocyte concentrates (buffy coats) obtained from the blood bank at Sahlgrenska University Hospital, Gothenburg, Sweden using peripheral blood from healthy human blood donors. According to the Swedish legislation section code 4§ 3p SFS 2003:460, no ethical approval is needed since the buffy coats are provided anonymously.

### 2.3. Isolation of human neutrophils

Neutrophils were isolated from buffy coats obtained from healthy blood donors as previously described [20, 21], In brief, the buffy coat was mixed with dextran, dissolved in physiological saline, to a final concentration of 1%, for sedimentation of the erythrocytes (1 x *g*, 20 min, room temperature). The supernatant, containing the leukocytes, was then separated on a Ficoll-Paque gradient by centrifugation (930 x *g*, 15 min, 4°C) to pellet the neutrophils. Any remaining erythrocytes in the neutrophil pellet were removed by hypotonic lysis after which the isolated neutrophils were washed twice in KRG without CaCl_2_ (220 x *g*, 10 min, 4°C) before analyzing the neutrophil concentration and purity on a Sysmex KX-21N Hematology Analyzer (Sysmex Corporation, Kobe, Japan). Isolated neutrophils were diluted in KRG to a concentration of 1x10^7^ neutrophils/mL and stored on ice until use on the same day as they had been isolated. The purity (mean ± SD) of the isolated neutrophils from the 10 different donors used in the study was 92% ± 4%.

### 2.4. Measurement of the increase in intracellular calcium ion concentration [Ca^2+^]_i_

Isolated neutrophils were pelleted by centrifugation (220 x *g*, 10 min, 4°C). Thereafter, the supernatant was removed, and the pellet was resuspended to a concentration of 2x10^7^ neutrophils/mL in CaCl_2_ free KRG with BSA (0.1% w/v) and loaded with Fura 2-AM (2 µg/mL, 30 min at room temperature in darkness). The cells were then washed twice; first with RPMI 1640 medium and then with KRG by centrifugation (300 x *g*, 10 min, 22°C). After aspiration of the supernatant the stained and washed neutrophil pellet was resuspend in KRG to a concentration of 2x10^7^ neutrophils/mL and stored on ice in darkness until analysis of [Ca^2+^]_i_. Measurements of [Ca^2+^]_i_ was performed on a Perkin Elmer fluorescence spectrophotometer (LC50B), with excitation wavelengths of 340 nm and 380 nm, an emission wavelength of 509 nm, and slit widths of 5 nm and 10 nm, respectively using 4.2 mL disposable polystyrene cuvettes and a 2.475 mL reaction mixture containing KRG and Fura-2 loaded neutrophils (5 x 10^6^). The reaction mixture was equilibrated for 10 minutes at 37°C, before addition of a stimulus (25 µL) and measurement of Fura-2 fluorescence continuously over time. For experiments where the effects of Larixol or YM-254890 were determined, these compounds were added to the reaction mixture just prior the ten minutes equilibration phase. The transient rise in [Ca^2+^]_i_ is presented as the ratio of fluorescence intensities (340 nm: 380 nm) detected. For analysis of peak [Ca^2+^]_i_, the peak value was subtracted by the background level, i.e., the value recorded just prior stimulation with the respective agonist.

### 2.5. Measurement of superoxide anions generated by the NADPH-oxidase

An isoluminol-enhanced chemiluminescence (CL) technique was used to measure the release of superoxide anions (the precursors of reactive oxygen species [ROS]) generated upon activation of the neutrophil NADPH-oxidase as described [22, 23]. The CL measurements were performed in a six-channel Biolumat LB 9505 (Berthold Co., Wildbad, Germany), using 4 mL disposable polypropylene tubes and a 900 µL reaction mixture containing KRG, isolated neutrophils (10^5^), isoluminol (2 x 10^-5^ M) and HRP (4 U/mL). The reaction mixture was equilibrated for five minutes at 37°C, before addition of a stimulus (100 µl) and measurement of light emission/super oxide anion production continuously over time. For experiments where the effects of Larixol or YM-254890 were determined, these compounds were added to the reaction mixture just prior to the five minutes equilibration phase. For analysis of peak superoxide anions produced, the background level, i.e., the value recorded just prior stimulation with the respective agonist was subtracted from the peak value by. The light emission/superoxide anion production is expressed as Mega counts per minute (Mcpm).

### 2.6 Data analysis

Data analysis was performed using GraphPad Prism 10 for macOS, version 10.0.0 (Graphpad Software, San Diego, CA, USA). Statistical analysis was performed using a paired Students *t*-test or a repeated measures one-way ANOVA followed Sidak’s multiple comparison test. The exact number of repeats performed independently on neutrophils isolated from different individuals (n) and the specific statistical test used for each figure are described in the figure legends. Statistically significant differences are indicated by \**p* < 0.05, \*\*\**p* < 0.001, \*\*\*\**p* < 0.0001.

## 3. Results

### 3.1. The Gα_q_ inhibitor YM-254890 lacks inhibitory effect on the FPR1 and FPR2 mediated rise in the intracellular concentration of free calcium ions ([Ca^2+^]_i_)

It is well established that the Gα subunit, dissociated from the βγ complex upon activation of a Gα_q_containing G protein, activates phospholipase C (PLC) that hydrolyzes phosphatidyl-inositol-4,5-bisphosphate (Ptdlns(4,5)P2/PIP_2_) to inositol trisphosphate (IP_3_) and diacylglycerol (DAG). When released to the cytosol, soluble IP_3_ then activates IP_3_-receptors on the Ca^2+^ storing endoplasmic reticulum, and this activation results in a release of Ca^2+^ from the storage organelles and by that an increase in the cytosolic concentration of free calcium ions ([Ca^2+^]_i_). Interestingly, the same signaling pathway is also activated by the dissociated βγ subunit down-stream of receptors that are coupled to Gα_i_containing G proteins [7]. In this study we investigated the [Ca^2+^]_i_ in human neutrophils upon activation of two Gα_i_coupled GPCRs (FPR1 and FPR2) and two Gα_q_ coupled GPCRs (PAFR and P2Y_2_R) by four receptor selective orthosteric agonists (fMLF, agonist for FPR1 [19], WKYMVM, agonist for FPR2 [24], PAF, agonist for the PAFR [25] and ATP, agonist for the P2Y_2_R [26, 27]) that all have in common that they can initiate a PLC dependent increase in [Ca^2+^]_i_ (Fig. 2A-B [7]). The rise in [Ca^2+^]_i_ down-stream of the PAFR and the P2Y_2_R was completely abolished in the presence of the tool compound YM-254890, a cyclic depsipeptide isolated from *Chromobacterium* sp. QS3666 shown to act as a selective Gα_q_ inhibitor ([15]). In contrast, the rise in [Ca^2+^]_i_ induced by the FPR agonists (fMLF and WKYMVM for FPR1 and FPR2, respectively) was not affected by the presence of YM-254890 (Fig 2A-B [7]). Hence, these data support the Gα_i_ coupling of signaling down-stream of FPRs (being YM-254890 insensitive and pertussis toxin sensitive [28, 29]).

**Figure 2.**
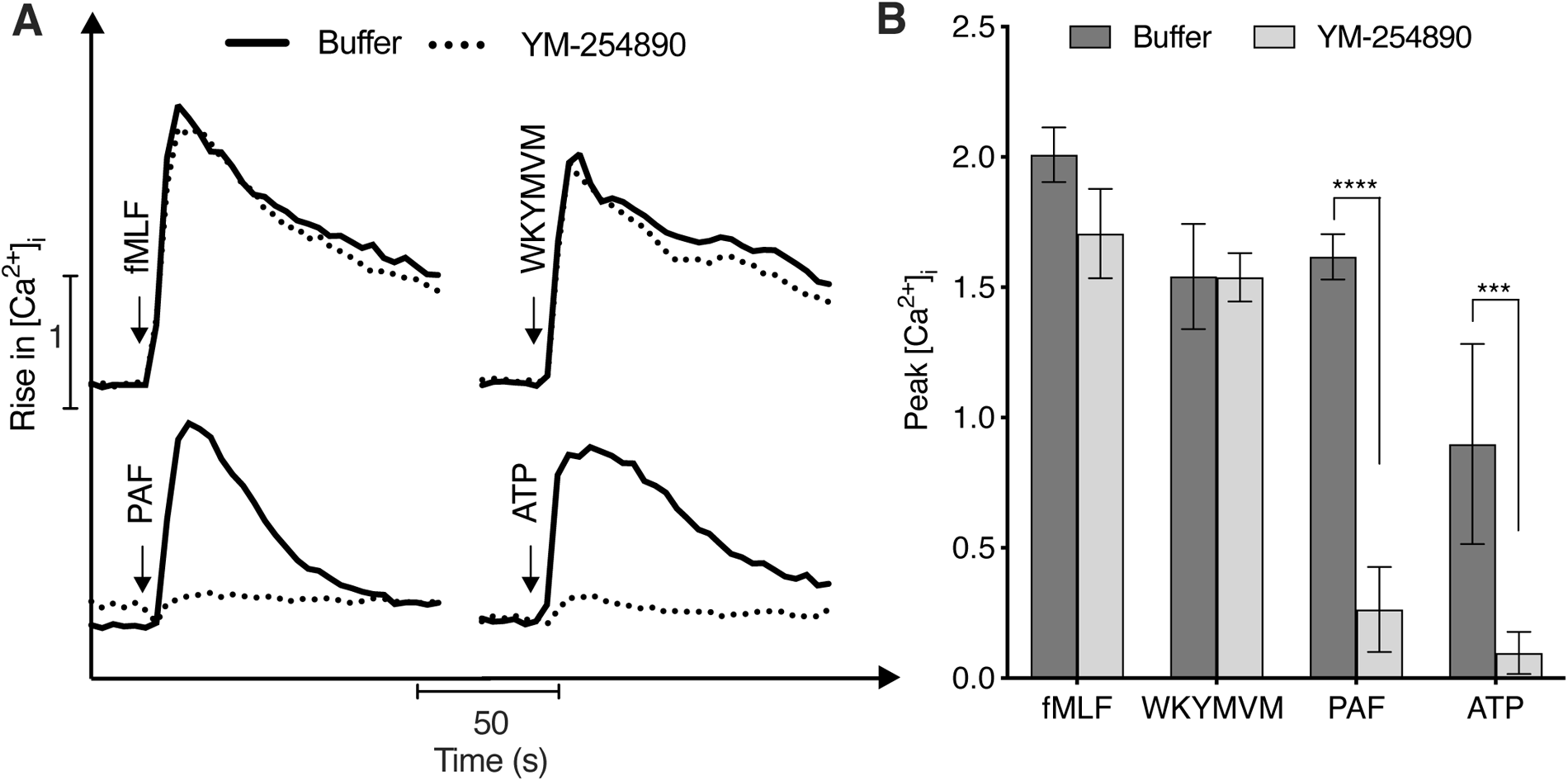
Gα_q_ inhibition does not block the FPR1 or FPR2 mediated rise in intracellular calcium concentrations [Ca^2+^]_i_. Fura-2 loaded neutrophils were incubated for ten min at 37 °C in the absence (buffer) or presence of the Gα_q_ inhibitor YM-254890 (200 nM) prior to stimulation with fMLF (10 nM), WKYMVM (10 nM), PAF (1 nM) or ATP (5 µM). **A**) One representative trace of the different agonists induced (time of stimulation is indicated by an arrow) transient rise in intracellular calcium ions [Ca^2+^]_i_ measured over time. **B**) Summary of the peak [Ca^2+^]_i_ (mean ± SD, n = 3). Statistically significant differences in **B** were evaluated by a repeated measures one-way ANOVA with Sidak’s multiple comparisons test for each agonist in the absence or presence of YM-254890 (\**p*<0.05, \*\**p*<0.01).

### 3.2. Larixol lacks inhibitory effect on the rise in the intracellular concentration of free calcium ions ([Ca^2+^]_i_) induced by activation of human neutrophil expressed FPRs

According to earlier published data, Larixol affects neutrophil functions in human neutrophils incubated with the compound (at concentrations ≥ 2 µM) for a minimum of three minutes at 37°C [17]. As the signals transmitted by βγ complexes of Gα_i_ containing G proteins [17] constitute the bases for inhibition, Larixol should be expected to be without effect on the rise in [Ca^2+^]_i_ mediated by an activation of the Gα_q_ coupled PAFR and the P2Y_2_R, using PAF or ATP, respectively, as activating agonists. Accordingly, Larixol was without effect on the ATP-induced rise in [Ca^2+^]_i_, but displayed a minor inhibitory effect on the rise in [Ca^2+^]_i_ when neutrophils were activated with PAF (Fig. 3A-B). Based on the suggested inhibitory mechanism of action for Larixol [17] the compound should be expected to inhibit not only the transient rise in [Ca^2+^]_i_ induced by fMLF (the prototype FPR1 agonist), but also by WKYMVM, an agonist targeting FPR2 which is closely related to FPR1 (69% amino acids similarity) and that also couple to a Gα_i_ containing G protein [18, 19]. However, our data show that Larixol lacks inhibitory effect on the rise in [Ca^2+^]_i_ induced in neutrophils activated by either fMLF or WKYMVM (Fig. 3A-B).

**Figure 3.**
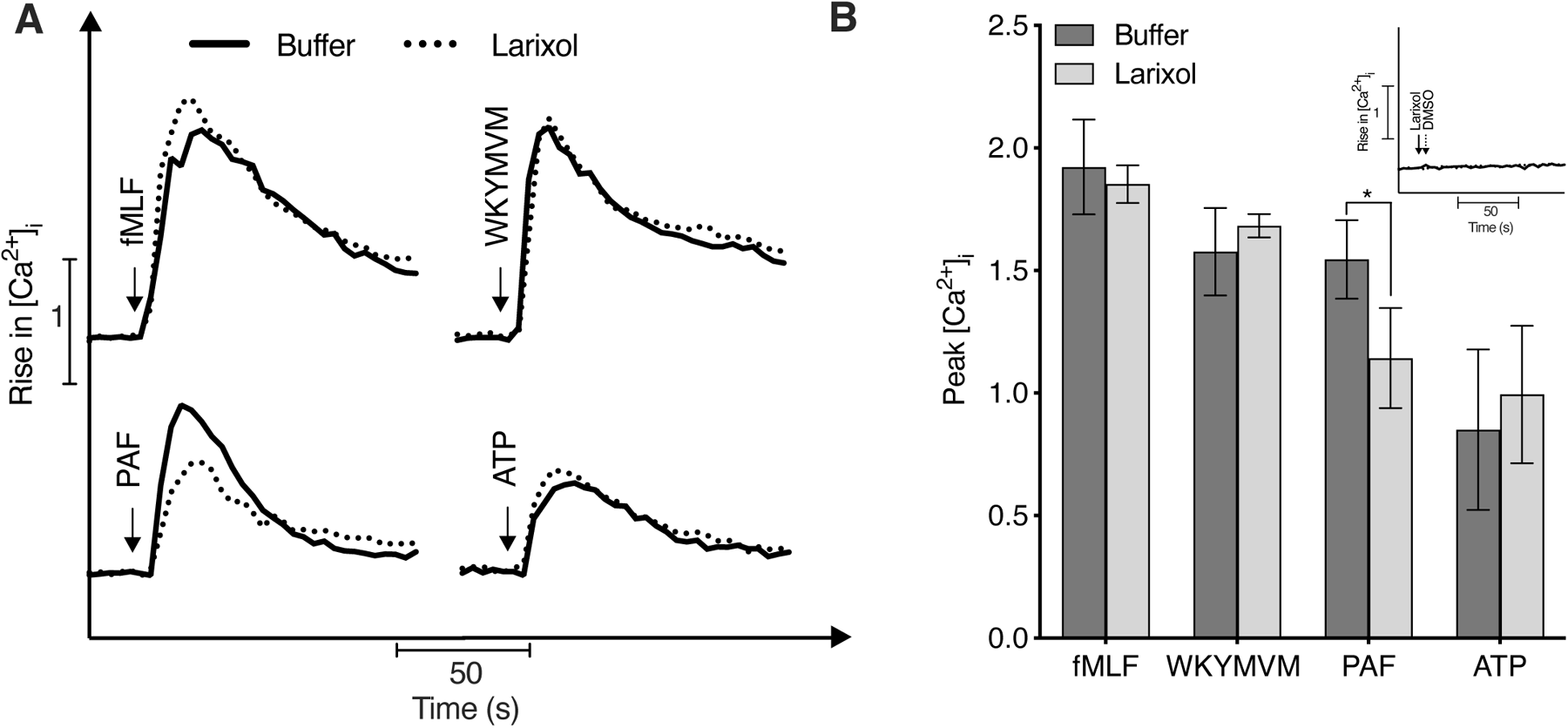
Synthetic Larixol lacks inhibitory effect on the FPR1 and FPR2 mediated rise in intracellular calcium concentrations [Ca^2+^]_i_. Fura-2 loaded neutrophils were incubated for ten min at 37 °C in the absence (buffer) or presence of Larixol (10 µM, synthesized by Biosynth) prior to stimulation with fMLF (10 nM), WKYMVM (10 nM), PAF (1 nM) or ATP (5 µM). **A**) One representative trace of the different agonists induced (time of stimulation is indicated by an arrow) transient rise in intracellular calcium ions [Ca^2+^]_i_ measured over time. **B**) Summary of the peak [Ca^2+^]_i_ (mean ± SD, n = 4). **Inset**) Fura-2 loaded neutrophils were incubated for ten min at 37 °C before stimulated (indicated by an arrow) with Larixol (10 µM, solid line) or KRG/DMSO as a vehicle control (0.001% v/v, dotted line). and measurement of [Ca^2+^]_i_ over time. One representative transient rise in [Ca^2+^]_i_ out of four individual experiments is shown. Statistically significant differences in **B** were evaluated by a repeated measures one-way ANOVA with Sidak’s multiple comparisons test for each agonist in the absence or presence of Larixol (\**p*<0.05).

These data thus demonstrate that in contrast to the described inhibitory effect of Larixol on the fMLF-induced rise in [Ca^2+^]_i_ [17] no such inhibitory effect is achieved using a synthetic variant of Larixol. Taken together, the data thus imply that Larixol lacks a general inhibitory effect on the down-stream signaling by Gα_i_ as well as Gα_q_ coupled neutrophil GPCRs.

### 3.3. Larixol lacks inhibitory effect on the FPR1 and FPR2 mediated superoxide anion production

According to earlier published data, Larixol should also inhibit the fMLF-induced activation of the superoxide anion producing NADPH-oxidase in human neutrophils [17]. As FPR agonist mediated activation of the NADPH-oxidase can occur without a concomitant rise in [Ca^2+^]_i_ [30, 31], we next investigated if Larixol displayed any inhibitory effect on the NADPH-oxidase activity mediated by the same agonists used to determine the effect of Larixol on the transient rise in [Ca^2+^]_i_. Three of these agonists (fMLF, WKYMVM and PAF), activate the superoxide oxide generating NADPH-oxidase in naïve neutrophils whereas the NADPH-activating signals generated by ATP (through P2Y_2_R), is blocked by the actin cytoskeleton in naïve neutrophils but can become activated in the presence of actin cytoskeleton disrupters [26]. However, like our calcium data (Fig. 3), but in contrast to earlier published data [17] Larixol lacked inhibitory effect not only on the NADPH-oxidase response induced by fMLF, but also on the response induced by WKYMVM, ATP and PAF (Fig. 4A-E). No NADPH-oxidase activity was induced when stimulating neutrophils with Larixol alone (Fig. 4F) and in accordance with the earlier published study [17], Larixol had no effect on the NADPH-oxidase activity induced by PMA (Fig. 4G-H), a phorbol ester that activates the NADPH-oxidase via a direct activating effect on protein kinase C, *i.e.,* independent of any neutrophil plasma-membrane expressed receptor. In summary, these data show that in contrast to what has been shown for Larixol extracted from the root of *Euphorbia formosana* [17], synthetic Larixol does not exhibit any inhibitory effect on the fMLF-induced NADPH-oxidase activity.

**Figure 4.**
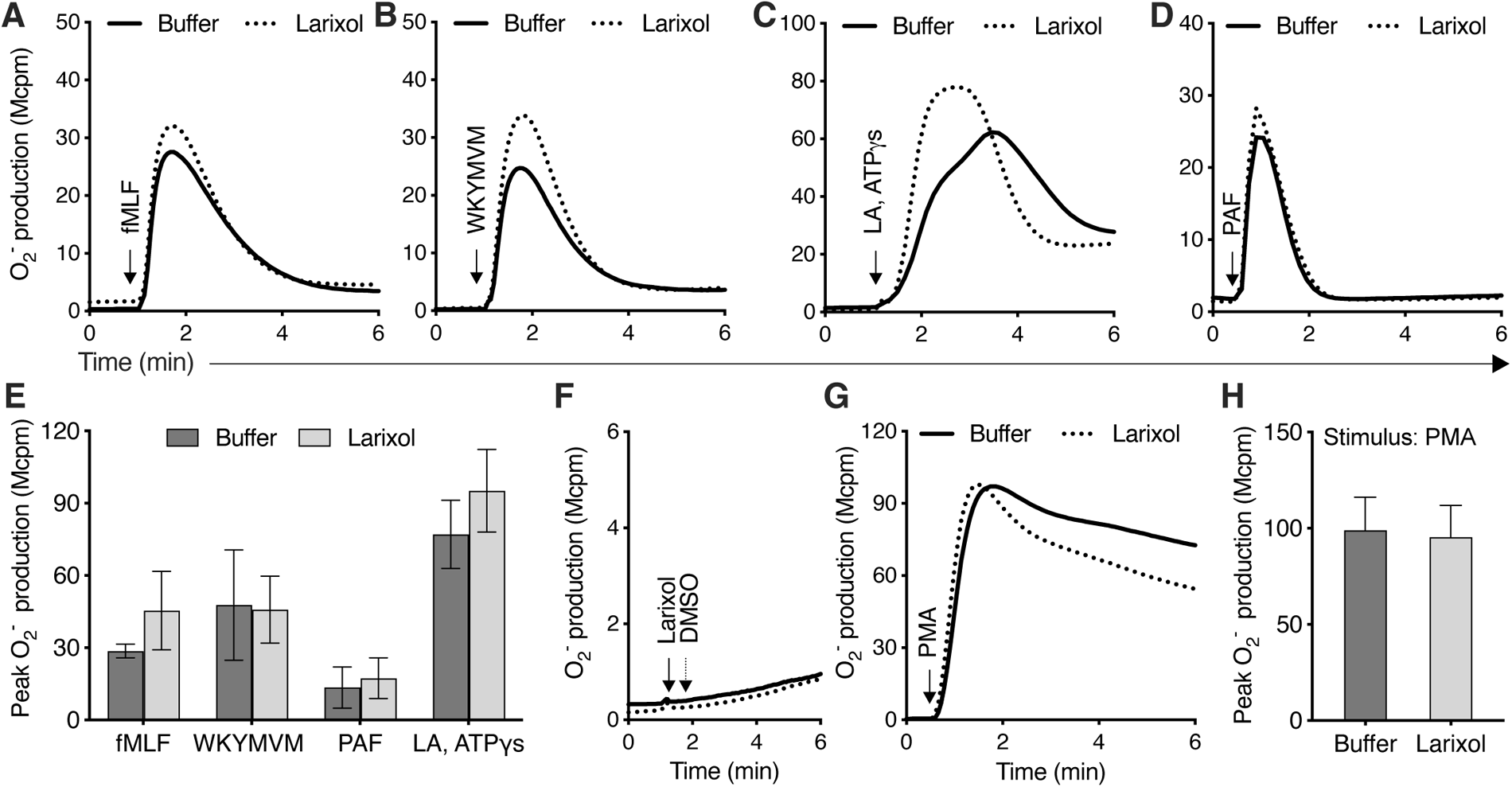
Synthetic Larixol lacks inhibitory effect on the FPR1 and FPR2 mediated NADPH-oxidase activity. Naïve human neutrophils were incubated at 37°C for five min in the absence (buffer) or presence of Larixol (10 µM, synthesized by Biosynth) prior stimulation with diverse agonists and measurement of NADPH-oxidase derived oxygen radical (O ^-^) production by isoluminol-amplified chemiluminescence over time. **A**-**E**) For activation of the NADPH-oxidase with ATPγs, the neutrophils were treated with the cytoskeleton disrupter Latrunculin A (LA, 25 ng/mL) during the five min at 37°C. One representative trace of the NADPH-oxidase superoxide anion (O ^-^) production upon stimulation (indicated with an arrow) with **A**) fMLF (100 nM), **B**) WKYMVM (100 nM), **C**) ATPγs (50 µM) or **D**) PAF (100 nM) is shown. **E**) Summary of the peak O ^-^ production (mean ± SD, n = 3). **F**) One representative trace out of three individual experiments is shown for neutrophils pre-incubated with buffer prior stimulation (indicated by an arrow) with Larixol (10 µM, solid line) or KRG/DMSO as a vehicle control (0.001% v/v, dotted line). **G**-**H**) One representative trace of the NADPH-oxidase superoxide anion (O ^-^) production upon stimulation (indicated with an arrow) with **G**) PMA (50 nM), summarized as the peak O ^-^ production in **H** (mean ± SD, n = 3). Statistically significant differences in **E** were evaluated by a repeated measures one-way ANOVA with Sidak’s multiple comparisons test for each agonist in the absence or presence of Larixol and in **H** by a paired Student’s *t*-test.

### 3.3. Natural Larixol lacks inhibitory effect on the FPR1 and FPR2 mediated superoxide anion production

The published results, showing that Larixol inhibits Gα_i_ protein signaling, were obtained using an extract from the roots of *Euphorbia formosana* [17]. Our results described above, showing that Larixol has no effect on GPCR signaling in neutrophils, were obtained with a synthetic compound. To determine the importance of the origin of the compound, we included a commercially available natural Larixol in our study. This natural Larixol was isolated from the turpentine oil of *Larix sp*. However, also this compound lacked inhibitory effect on the human neutrophil NADPH-oxidase activity induced by either of the four agonists (fMLF, WKYMVM, PAF and ATP) included in the study (Fig. 5A-E) but displayed a minor potentiated NADPH-oxidase activity upon activation with ATP. No NADPH-oxidase activity was induced by Larixol alone (Fig. 5F) and the compound was without effect also on the PMA-induced NADPH-oxidase activity (Fig. 5G-H).

**Figure 5.**
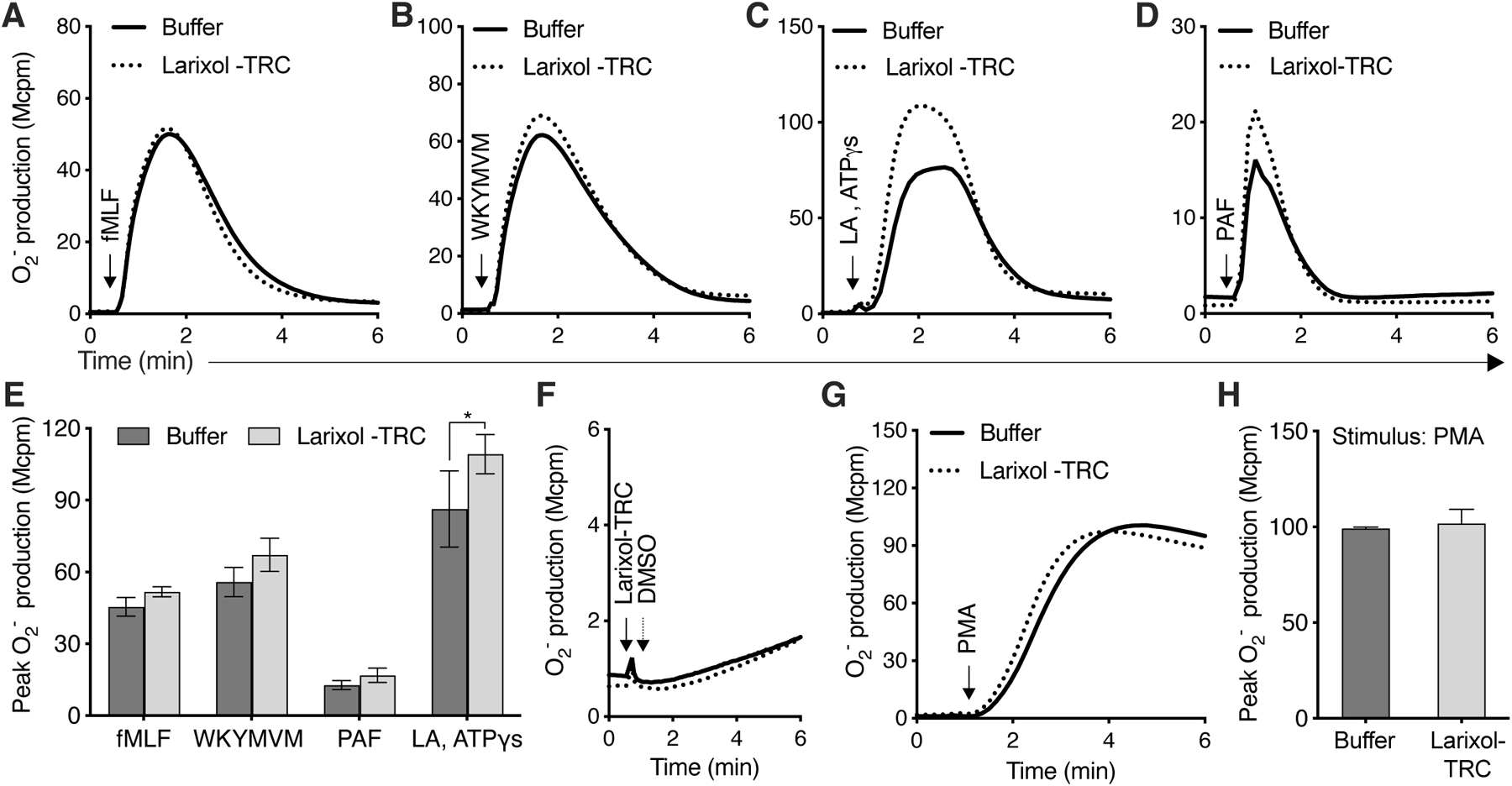
Natural Larixol, isolated from turpentine oil of *Larix sp*, lacks inhibitory effect on the FPR1 and FPR2 mediated NADPH-oxidase activity. Naïve human neutrophils were incubated at 37°C for five min in the absence (buffer) or presence of natural Larixol (10 µM, isolated from turpentine oil of *Larix sp by* Toronto research chemicals [TRC]; Larixol-TRC) prior to stimulation with diverse agonists and measurement of NADPH-oxidase derived oxygen radical (O ^-^) production by isoluminol-amplified chemiluminescence over time. **A**-**E**) For activation of the NADPH-oxidase with ATPγs, the neutrophils were treated with the cytoskeleton disrupter Latrunculin A (LA, 25 ng/mL) during the five min at 37°C. One representative trace of the NADPH-oxidase superoxide anion (O ^-^) production upon stimulation (indicated with an arrow) with **A**) fMLF (100 nM), **B**) WKYMVM (100 nM), **C**) ATPγs (50 µM) or **D**) PAF (100 nM) is shown. **E**) Summary of the peak O ^-^ production (mean ± SD, n = 3). **F**) One representative trace out of three individual experiments is shown for neutrophils pre-incubated with buffer prior stimulation (indicated by an arrow) with Larixol (10 µM) or KRG/DMSO (0.001% v/v, used as a vehicle control). **G**-**H**) One representative trace of the NADPH-oxidase superoxide anion (O ^-^) production upon stimulation (indicated with an arrow) with **G**) PMA (50 nM), summarized as the peak O ^-^ production in **H** (mean ± SD, n = 3). Statistically significant differences in **E** were evaluated by a repeated measures one-way ANOVA with Sidak’s multiple comparisons test for each agonist in the absence or presence of Larixol and in **H** by a paired Student’s *t*-test.

Taken together these data show that commercially available Larixol, synthesized or natural isolated from the turpentine oil of *Larix sp*, does not display any inhibitory effect on fMLF-induced neutrophil functions.

## 4. Discussion

A recent study identified Larixol, a diterpene extracted from the root of *Euphorbia formosana*, as a novel inhibitor of human neutrophil functions induced by fMLF, an orthosteric agonist for the Gα_i_ coupled GPCR termed FPR1. The mechanism of action for this inhibitory effect was suggested to occur by blockage of G protein signaling through *i)* a direct interaction between the activated βγ complex associated with the Gα_i_ subunit that together comprise the heterotrimeric G protein coupled FPR1 and *ii)* an indirect effect on selected FPR1 down-stream signaling kinases and enzymes [17]. In this study, we set out to characterize the GPCR selectivity of the inhibitory effects of Larixol in human neutrophils and to analyze the potential of the compound to act as a novel inhibitor of Gα_i_coupled human neutrophil expressed GPCRs in general. To do this we applied two commercially available variants of Larixol, one synthetic variant and one variant isolated from the turpentine oil of *Larix sp*. We then followed the methods described in the previous study [17] and pre-incubated isolated human neutrophils with Larixol (≥ 2 µM, for a minimum of three minutes at 37°C) prior to stimulation with different GPCR selective agonists and analysis of Larixol’s inhibitory effect on the transient rise in intracellular calcium ions and production of NADPH-oxidase derived superoxide anions. For thorough characterization of Larixol’s inhibitory effect we specifically choose four agonists that either selectively activates two different human neutrophil expressed Gα_i_ coupled GPCRs or two different human neutrophil expressed Gα_q_ coupled GPCRs [7]. Hence, the peptides fMLF and WKYMVM were used for activation of the Gα_i_ coupled (pertussis toxin sensitive) FPR1 and FPR2, respectively [28, 29], whereas PAF and ATP were used for activation of the Gα_q_ coupled (YM-254890 sensitive) PAFR and the P2Y_2_R, respectively (this study and [32, 33]). However, despite incubating neutrophils with up to 10 µM Larixol for ten minutes at 37°C prior to stimulation (the highest concentration and longest incubation time described for inhibition of fMLF mediated responses in the previous study [17]) and stimulating neutrophils with a lower concentration of fMLF for the measurement of the transient rise in intracellular calcium ions (10 nM, instead of 100 nM as used in the previous study [17]) our data did not reveal any inhibitory effect on the fMLF-induced human neutrophil responses. Furthermore, as our data also showed that Larixol was without effect on the human neutrophil responses induced by WKYMVM, our results strongly imply that Larixol is not a general inhibitor of Gα_i_ coupled human neutrophil GPCRs. Moreover, our results also imply that Larixol is not a general inhibitor of Gα_q_ coupled neutrophil expressed GPCRs, as the compound only had a minor inhibitory effect on the increase in intracellular calcium ions when induced by PAF but not by ATP and as Larixol had no effect on the NADPH-oxidase activity induced by PAF but showed a minor potentiating effect of this response when induced by ATP.

That is, we could not repeat the data by Liao *et al* showing an inhibitory effect on fMLF-induced neutrophil responses when using Larixol obtained as a diterpene extract from the root of *Euphorbia formosana*, a flowering plant that is native on a chain of islands southwest of Japan down to Taiwan. Based on the fact that the two commercially available sources of Larixol used in this study had a solvent purity > 95 % and lacked effects on the fMLF-induced neutrophil signaling/function, it is reasonable to assume that the extract of Larixol used by Liao *et al* not only contains Larixol but also other compounds that might be responsible for the demonstrated inhibitory effects in their study [17]. However, this is hard to decipher as *i)* the exact purity of Larixol in the extract used by Liao *et al* is not specified, *ii)* the researchers responsible for the study do not have the extract used in the study in sufficient amounts to provide to other researchers and *iii)* as we have not been able to find a commercial source of Larixol extracted from the root of *Euphorbia formosana*. It should be noted that some of the researchers involved in the work that described Larixol as an inhibitor of fMLF-induced neutrophil functions [17] also has shown that fMLF-induced neutrophil functions can be inhibited by another compound (2’,3-dihydroxy-5-methoxybiphenyl [RIR-2]) extracted from the root of the Asian plant termed *Rhaphiolepis indica* (L.) Lindl. *ex* Ker var*. tashiroi* Hayata *ex* Matsum. & Hayata (Rosaceae) [34]. Similar to the described mechanism of action for Larixol [17], the inhibitory effect of RIR-2 was suggested to be due to interference with/inhibition of signals generated by parts of the βγ complexes of the Gα_i_ containing G proteins coupled to FPR1 [34]. Yet, it awaits to be shown if commercially available RIR-2 demonstrates a similar inhibitory effect.

The most commonly used and best-established inhibitor to date for determination of if a GPCR is coupled to Gα_i_ is pertussis toxin (an exotoxin produced by the bacterium *Bordetella pertussis*) [9]. Pertussis toxin comprises six subunits, one pentameric cell-binding subunit and one catalytic subunit and is taken up by cells by receptor-mediated endocytosis upon binding to sialylated glucoconjugates on the plasma membrane. The exact endosomal route of pertussis toxin from the endosome to the cytosol is not known, but well in the cytosol pertussis toxin inactivates Gα_i_ subunits through ADP-ribosylation resulting in that Gα_i_ coupled GPCRs uncouples from the Gα_i_ subunit and thereby the signal transduction down-stream the receptor becomes disrupted [35]. However, determination of if a GPCR is coupled to Gα_i_ using pertussis toxin has certain limitations including some Gα_i_ subunit independent effects such as enhancement of immune responses (summarized in [12]) and doubtful specificity for inhibition of only the Gα_i_ subunit and not the Gα_q_ subunit regarding GPCRs expressed by human neutrophils [7]. We and others have previously shown that, as expected, treatment of human neutrophils with pertussis toxin for ≥ 1.5 hours inhibits signaling/functions mediated by activation of FPR1 or FPR2 with fMLF or WKYMVM, respectively [28, 32, 36]. However, when it comes to human neutrophils the G protein selectivity of pertussis toxin has been put into question as pertussis toxin treatment also induce inhibition of the Gα_q_ coupled PAFR [13, 14]. The large similarities between the G proteins makes it hard to design/find inhibitors that selectively blocks signaling down-stream of receptors expressed in naïve cells. Although research to find new inhibitors is ongoing [37], the tools that are currently available are limited and every attempt to find new tool compounds is much appreciated by the research community. However, the results in this study together with our previous results in which we showed that a presumed Gα_q_ inhibiting pepducin (P4Pal_10_, shown to inhibit responses mediated by several cell line expressed Gα_q_-coupled GPCRs [38]), does not inhibit Gα_q_-coupled GPCRs (PAFR and/or P2Y_2_R) expressed by human neutrophils and monocytes but instead acts as a selective inhibitor of Gα_i_-coupled FPR2 mediated responses in these [39] clearly show that investigations of the precise mechanism and selectivity for a G protein as well as for selected GPCRs are required when new tool compounds targeting a specific Gα subunit are introduced.

The fact that specific Gα_q_ inhibitors recently have become available and demonstrated as very useful tool compounds for determination of if a GPCR is coupled to Gα_q_, has reduced the need for new tool compounds for Gα_q_ selective compounds [5, 7, 15, 16].

As the Gα_i_subunit family has been reported to play an important role in several immune system functions and comprise the largest and most diverse class of the four Gα subunit classes (Gα_s_, Gα_12/13_, Gα_q_ or Gα_i_) and [40], specific inhibitors of Gα_i_ containing G proteins are eagerly awaited. The exact pharmacological approach for design of such inhibitors has several challenging obstacles (summarized in [9]) but the results presented in this study strongly implies that Larixol can be ruled out in the search for novel, potent and selective Gα_i_ inhibitors.

## Abbreviations

BSA: bovine serum albumin
DMSO: dimethyl sulfoxide
fMLF: *N*-formyl-methionyl-leucyl-phenylalanine
FPR: formyl peptide receptor
GPCR: G protein-coupled receptor
HRP: horseradish peroxidase
KRG: Krebs-Ringer Glucose phosphate buffer
Mcpm: Mega counts per minute
NADPH: nicotinamide adenine dinucleotide phosphate hydrogen
PAF: platelet activating factor
PMA: phorbol 12-myristate 13-acetate
WKYMVM: tryptophyl-lysyl-tyrosyl-methionyl-valyl-methioninamide
[Ca^2+^]: intracellular calcium ion concentration

## CRediT author statement

**Lena Björkman**; funding acquisition, investigation, writing – review & editing. **Huamei Forsman**; conceptualization, funding acquisition, methodology, resources, validation, writing – review & editing. **Linda Bergqvist**; data curation, investigation, writing – review & editing. **Claes Dahlgren**; conceptualization, methodology, resources, supervision, validation, writing – original draft, writing – review & editing. **Martina Sundqvist**; conceptualization, data curation, formal analysis, funding acquisition, investigation, methodology, project administration, supervision, validation, visualization, writing – original draft, writing – review & editing.

## Declaration of Competing Interest

The authors declare that they have no known competing financial interests or personal relationships that could have appeared to influence the work reported in this paper.

## Data availability

Data will be made available on request.

## Acknowledgment

The work was financed by the Swedish Medical Research Council (HF; 2018-02848 and 2022-00624) and by grants from the Swedish state under the agreement between the Swedish government and the country councils, the ALF-agreement (HF; ALFGBG78150), the Clas Groschinskys Memorial Fund (MS; M21146), the Åke Wiberg Foundation (MS; M21-0025), the Sahlgrenska International Starting Grant (MS; GU2021/1070), the Magnus Bergwall Foundation (MS; 2021-04110), the King Gustaf V 80-Year Foundation (HF: FAI-202-0687; MS; FAI-2021-0804 and FAI-2022-0873) the Rune and Ulla Amlövs Foundation (MS: 2023-418; LB:2021-312), and Reumatikerförbundet (LB: R-941156 and R-969625).

## Notes

### Competing Interest Statement

The authors have declared no competing interest.

